# Unilateral neglect is associated with poor proprioception after stroke – a systematic review

**DOI:** 10.1101/710921

**Authors:** Georgia Fisher, Camila Quel de Oliveira, Simon Gandevia, David Kennedy

## Abstract

**Background:** Proprioceptive impairment is a potential contributing factor to the clinical presentation of Unilateral Neglect (UN), a common and debilitating condition that can occur after stroke. To date there has not been a comprehensive review of studies examining the various aspects of proprioception in UN after stroke.

**Aim:** To determine if the presence of UN is associated with more severe proprioceptive deficit in stroke affected populations.

**Methods:** The MEDLINE, Embase, Scopus, CINAHL and Web of Science databases were searched from inception to January 2019 using an a priori search strategy. Two independent reviewers screened abstracts and full texts. Two reviewers then independently extracted data from each full text. A third reviewer resolved disagreements at each step. Risk of bias was assessed using the AXIS Quality Assessment tool. For full protocol see PROSPERO, registration number CRD42018086070. One-hundred and sixty-seven abstracts were identified, of which fifty-four were eligible for full text screening. A total of 18 papers were included in the review.

**Conclusions:** More severe proprioceptive deficit is associated with the presence of UN after stroke. However, the available evidence is limited by the large heterogeneity of assessment of both UN and proprioception, and level of study quality. UN and proprioception are seldom completely assessed in research, and it is likely this is true in everyday clinical practice.

## Background

Unilateral neglect (UN) is a complex condition that can result from a stroke, characterised by the failure to report, respond, or orient to novel or meaningful stimuli that are presented on the side opposite the brain lesion [1]. UN can impact on multiple spaces and modalities [2, 3], and is associated with right brain lesions, larger lesion volumes, and advanced age [4-6]. UN is linked to greater length of hospital stay, higher incidence of falls, poorer functional outcome, and a reduced likelihood of home as a discharge location [7-11]. Between 30% and 60% of patients with UN remain functionally impaired a year post-stroke [12-14]. General estimates of the incidence of UN after stroke range from 23.5% to 67.8% [4, 7].

Despite the high incidence and negative functional consequences of UN, there is no consensus about effective, yet clinically feasible, assessment and treatment for the condition [15-19]. There are upwards of 28 different standardised assessments of UN, most of which only capture a single, non-functional domain of the condition [15]. Thus, it is likely that current assessment of UN does not provide a complete picture of functionally relevant patient impairments. This precludes the development and implementation of targeted treatment strategies and may explain the small to moderate effect sizes reported in reviews of UN treatment [18, 20, 21].

Reviews of UN report an association with poor motor recovery after stroke [22-24]. A critical contributor to the ability to control movement during functional tasks is proprioception [25-27]. Proprioception refers to a set of sensorimotor processes that enable the ability to detect movement and positions at different joints, judge forces exerted by muscles, time muscular contractions and develop knowledge of body representation [25]. Proprioception has been identified as an important factor in functional outcome after stroke [28-30]. Thus, more severe proprioceptive impairment is a potential contributing factor to the poor clinical outcomes associated with UN.

Proprioception can be conceptualised into ‘low-level’ or ‘high-level’ processes. Reporting whether your thumb has been moved up or down is an example of ‘low-level’ proprioception. This tests the integrity of peripheral receptors and early components of afferent processing. An example of ‘high-level’ proprioception is knowing where your arm is in relation to yourself and the world. Thus, ‘high-level’ proprioception refers to the integration of ‘low-level’ signals into internal models of the body, peri-personal space, and the world. While it may be useful to conceptually separate ‘low-level’ and ‘high-level’ proprioception, it is the combination of both that allow a complete realisation of the proprioceptive sense.

A recent review of UN and upper limb function identified a lack of research in sensorimotor interventions for the affected upper limb in patients with UN [22]. However, in order to develop these interventions, it is important to first identify the specific underlying sensorimotor impairments driving poor outcomes in this population. Proprioception represents one such impairment, although to date there has not been a comprehensive review of studies examining the various aspects of proprioception in UN after stroke. Hence, the aim of this systematic review is to determine if the presence of UN is associated with more severe proprioceptive deficit in stroke affected populations.

## Methods

The protocol for this systematic review is registered in PROSPERO, under the registration number CRD42018086070. It can be accessed at the following link: https://www.crd.york.ac.uk/prospero/display_record.php?RecordID=86070. The CINAHL, Embase, MEDLINE, Scopus, and Web of Science electronic databases were searched from inception to January 15, 2019 using an a priori strategy of search terms designed to capture all aspects of unilateral neglect and proprioception in a stroke population (S1 File). Article titles and abstracts were screened by two independent reviewers (G.F., D.K.) according to the following inclusion criteria: (1) adult participants aged 18-80 (2) first time stroke confirmed on medical imaging (3) included at least one standardised assessment of UN, and of proprioception and (4) outcome measures reported for patient groups with and without UN. There was no restriction on publication year. Baseline data from randomised controlled trials were also included provided that the full data set was available, and the study data were subsequently considered cross-sectional. Studies that used clinical tests that assessed only balance and/or vestibular function and/or motor function that was not specific to proprioception were excluded.

Full article texts of relevant articles were then retrieved and screened by the same reviewers. Conference proceedings and dissertations were not included. Authors were contacted to request full text or study data where it was not available. Studies in languages other than English were included and translated using an online translation service. A third reviewer resolved disagreements at each step of study selection (C.Q.).

Two reviewers (G.F., C.Q.) then independently extracted data from included studies using a standardised form based on the Cochrane Data Extraction Template [31]. Extracted information included: study aims, study setting, study population, participant demographics and baseline characteristics, study methodology, recruitment and study completion rates, outcomes measurements and their time points of collection, and the suggested mechanisms of interaction between proprioception and neglect. Study quality was evaluated using the Appraisal tool for Cross-Sectional Studies (AXIS), a 20 item scale developed using a Delphi panel consensus [32]. Disagreements were recorded and resolved through discussion and when necessary with a third author (D.K.). The AXIS acknowledges the issues with the summation of checklists for study quality [33, 34], and as such does not have published cut-off scores to categorise studies as low, medium, or high quality [32]. A descriptive synthesis of the findings stratified according to different categories of proprioception assessment was made. Where the necessary data were available, narrative descriptions were completed for people with different forms of neglect delineated according to modality (spatial, motor, or representational) and space affected (personal, peri-personal, or extra-personal).

## Results

One-hundred and sixty-seven abstracts were identified, of which fifty-four were eligible for full text screening. A total of eighteen papers were included in the review [35-52]. Figure 1 describes the selection process of the studies. The predominant reasons for exclusion at full text review were inadequate data reporting and a lack of a measure of proprioception. The full list of excluded studies and the reasons for their exclusion can be found in S2 Table.

**Fig 1.**
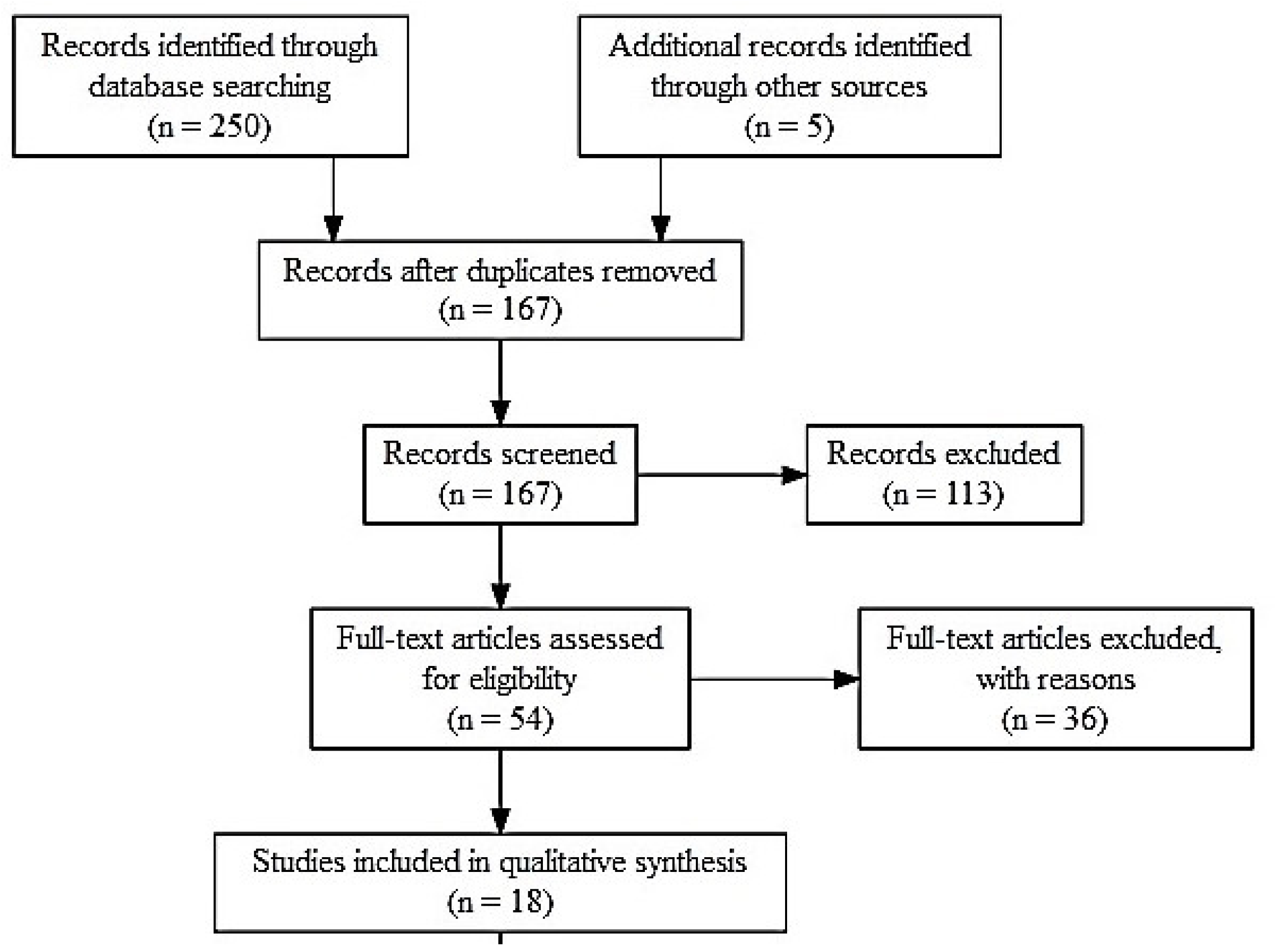
Selection Process Flowchart.

### Characteristics of Included Studies

The complete characteristics of the eighteen studies included are summarised in Table 1. The mean age was 60.3 ± 5.4 years, (UN+ = 60.9 ± 5.6, UN- = 59.8 ± 5.4) and the majority of participants were male (65%). Most studies recruited participants in the sub-acute phase (three weeks to six months post stroke), however two studies [44, 48] collected data exclusively from participants in the chronic phase (more than six months after stroke). Five studies [37, 39, 49-51] recruited mixed populations, and a single study [52] limited recruitment to the acute phase (less than three weeks post stroke).

**Table 1:**
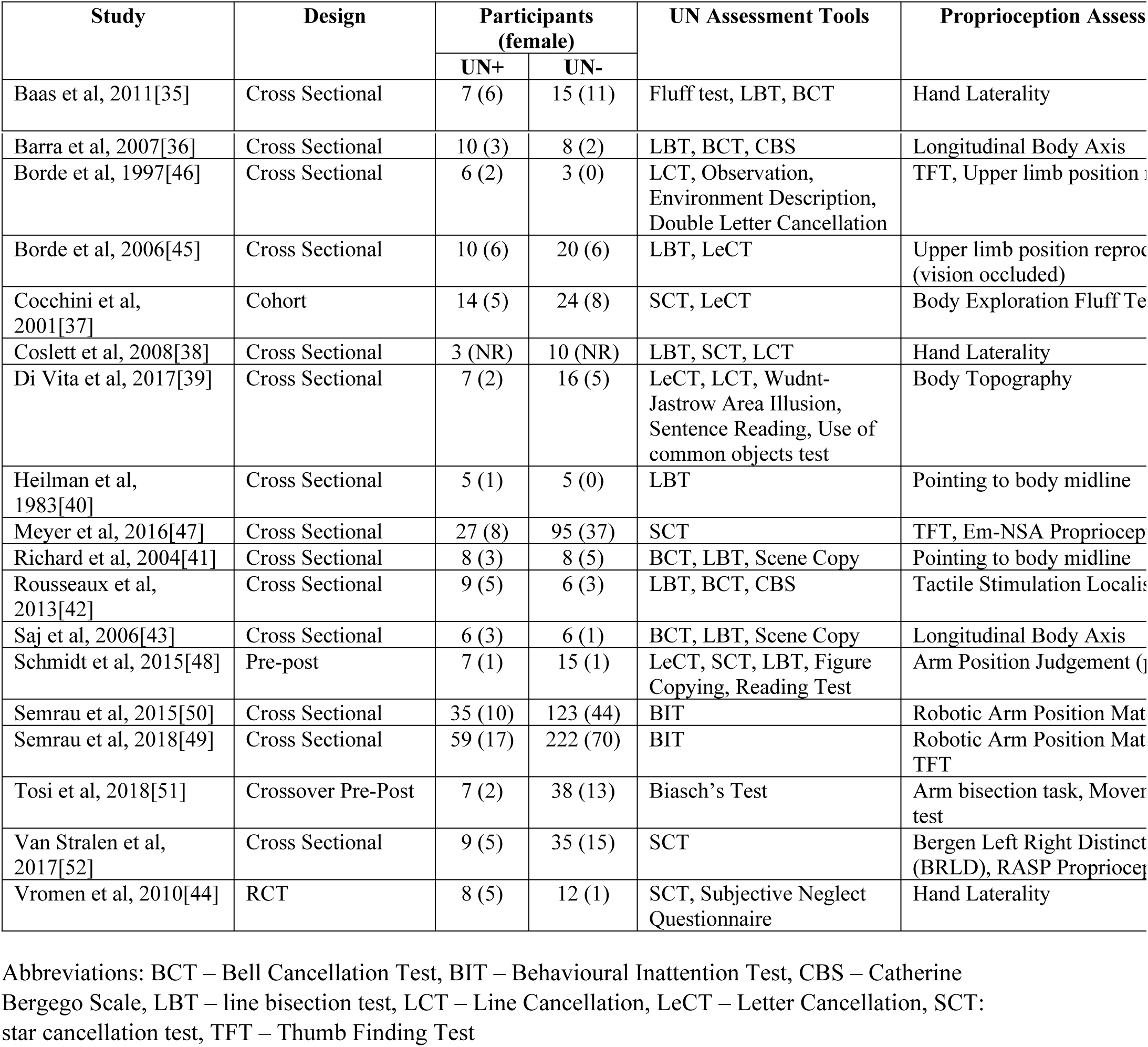
Included Study Descriptions.

There was a total of eighteen different assessment tools used to identify UN in the studies of this review. UN was assessed with tasks sensitive only to extra-personal UN in seven studies [37, 38, 40, 45, 46, 52, 53], and hence hemispatial UN is the predominant form of UN described. Two studies [35, 39] report using an assessment tool designed to capture personal neglect in addition to an extra-personal measure, one paper [51] reported use of a personal UN test alone, and two papers [49, 50] reported using a behavioural assessment in isolation. Two papers [36, 42] added a functional assessment of UN, the Catherine Bergego Scale (CBS).

This review found that proprioception was assessed by thirteen different methods. Three studies used a proprioceptive assessment restricted to detection or discrimination of movement [47, 48, 52] that can be considered ‘low level’. The remaining studies of this review assessed ‘high level’ aspects of proprioception. Four studies [45, 46, 49, 50] reported outcomes for proprioception measured with a limb matching task that required integration of motor planning. Of the four studies [35, 38, 44, 52] examining laterality, three [35, 38, 44] used a hand laterality task and one [52] utilised a laterality task in which participants were required to point to the left or right hand of human stick figure drawings in various orientations. Five studies [36, 40, 41, 43, 51] used an assessment that required participants to identify the location of the body midline, or a body axis. Finally, three studies [37, 39, 42] examined participant perception of body topography.

The results of the methodological quality assessment using the AXIS tool are summarised in Table 1, and fully reported in S3 Table. All studies defined their target population, reported internally consistent results, and justified their discussion and conclusions. However, there were multiple issues with quality across most of the studies included in this review. Only a single study [36] justified their sample size, and was the only study to report a method of measuring non-response to recruitment. Twelve papers failed to discuss limitations to their study [35, 37-43, 45, 46, 50, 51], and four studies did not use a previously validated assessment tool [37, 42, 45, 46]. Five studies did not present results for all planned analyses [37-41], and six were deficient in their description of basic participant demographic data [40, 42-44, 48, 52]. Finally, seven studies did not use consecutive patient selection, instead using convenience samples [35, 37, 40, 42, 43, 48, 50, 52].

### Proprioception Outcomes

Table 2 displays the results of each selected study stratified by type of proprioceptive assessment, including a description of the study outcomes, and study findings specific to this review. A total of fifteen studies reported more severe proprioceptive deficits in participants with UN after stroke compared to participants without the condition. Three studies reported no association between UN and more impaired proprioception, and a single study did not report sufficient data for judgement to be made.

**Table 2:** Study Result Summary.

| Study | Main Findings | Findings Relevant to Current Review | UN Associated with Proprioception Impairment? |
| --- | --- | --- | --- |
| **Movement Detection / Judgement** |  |  |  |
| Meyer *et al.* 2016 [47] | UN+ had significantly more frequent, and more severe somatosensory deficits compared to UN-. Proprioceptive deficits were significantly more frequent in UN+. | UN is associated with impaired movement detection, and body representation | Y |
| Schmidt *et al.*, 2013 [48] | Arm position sense was significantly more impaired in the left (contralesional) but not the right arm in RHD UN+ patients as compared with RHD UN- patients. Left-GVS selectively improved the accuracy of APS in RHD UN+ patients during stimulation, and 20 minutes after stimulation. | UN is associated with impaired passive arm position judgement | Y |
| van Stralen *et al.*, 2017 [52] | 100% of UN+ participants were impaired on movement detection task, compared to 26% of UN-. | Associated between UN and deficits not reported | UTD |
| **Joint Position Matching** |  |  |  |
| Borde *et al.*, 1997 [46] | Somatosensory disorders may be associated with deafferentation, motor impairment, or a non-specific cognitive disorder. No firm conclusions able to be drawn. | UN did not impact on position matching ability | N |
| Borde *et al.*, 2006 [45] | RHD participants significantly worse at reproducing passively imposed arm gestures than those with LHD. Defective RHD performance is independent of UN and constructive apraxia. | UN did not impact on position matching ability in RHD | N |
| Semrau *et al.*, 2015 [50] | 100% of UN+ failed the kinaesthetic task, compared to 59% of UN-. UN is | UN is associated with arm position | Y |
|  | predictive of the presence of kinesthetic deficits, however, kinesthetic deficits do not always indicate UN. | matching task failure, and altered kinematics |  |
| Semrau *et al.*, 2018 [49] | 67% percent of stroke affected participants had kinaesthetic impairments in the No Vision condition and 50% had kinesthetic impairments in the Vision condition. UN+ had larger impairments in both conditions compared to UN-. | UN is associated with arm position matching deficits, with and without the use of vision. | Y |
| **Laterality** |  |  |  |
| Baas *et al.*, 2011 [35] | UN+ had significantly worse scores on laterality tasks compared to UN-, indicative of a general dysfunction in contralesional representations in UN | UN+ was associated with impaired body representation | Y |
| Coslett, 1998 [38] | UN+ significantly worse than UN- at identifying pictures of left hands, indicating a disruption of or failure to access the left portion of the body schema | UN was associated with impaired body representation | Y |
| van Stralen *et al.*, 2017 [52] | Laterality ability is dissociable dependent on the perspective of the stimulus presented. Switching between first and third person perspective was predicted by proprioceptive impairments, attention and working memory. UN and finger agnosia predicted performance on laterality task performance. | UN associated with deficit in body representation | Y |
| Vromen *et al.*, 2011 [44] | UN+ scored significantly worse than UN- on the visual mental rotation task, but not on the mental rotation task with verbal instruction. | UN associated with impaired body representation | Y |
| **Body Axis / Midline** |  |  |  |
| Barra *et al.*, 2007 [36] | Abnormal LBA perception is associated with sensory loss, UN, and postural and gait disturbances in patients with stroke | UN+ was correlated with impaired body representation | Y |
| Heilman *et al.*, 1983 [40] | UN+ significantly worse on pointing to body midline task, which may be due to disruption to representational map, or hemispatial hypokinesia. | UN is associated with impaired body representation | Y |
| Richard *et al.*, 2004 [41] | UN+ displayed a significant rightward shift of body centred SSA compared to UN-. There was a significant correlation between body centred long line bisection and SSA in UN+ only. | UN is associated with an impairment in body representation | Y |
| Saj *et al.*, 2006 [43] | UN+ had a subjective counter clockwise tilt, and rightward translation of the perceived body midline, UN- not significantly deviated from midline. | UN is associated with impairment in body representation | Y |
| Tosi *et al.*, 2017 [51] | Mirror box training has a significant positive effect on arm bisection accuracy | There is no association between | N |
|  | in stroke participants. UN had no effect on arm bisection accuracy. | UN and body representation |  |
| **Body Topography** |  |  |  |
| Cocchini *et al.*, 2001 [37] | UN+ were significantly worse at detaching contralesional targets on the body, implicating a disruption in body representation. | UN was associated with impaired body representation | Y |
| Di Vita *et al.*, 2017 [39] | Personal UN+ were significantly worse at rearranging tiles of body parts. Extrapersonal UN has no impact on results. | Personal UN only is associated with impairment in body representation. | Y |
| Rousseaux *et al.*, 2013 [42] | UN+ had significant deviations in localisation of trunk landmarks compared to UN-, and an underestimation of the width on the body that was greatest on the contralesional side | UN is associated with complex reorganisation of body representation | Y |

Abbreviations: LBA = longitudinal body axis, N = no, UTD = unable to determine, Y = yes

### Personal UN vs Extra-personal UN

Two studies, Di Vita et al. [39] and Baas et al. [35], assessed personal neglect in addition to extra-personal neglect. The studies evaluated proprioception with measures of body representation (body topography and hand laterality). Both found significant differences in body representation between those with personal UN and those without, and that extra-personal UN (assessed with cancellation tasks) had no influence on proprioceptive task performance.

### Behavioural Unilateral Neglect

The Behavioural Inattention Test (BIT) was used to assess UN in two studies by Semrau and colleagues [49, 50]. Each assessed proprioception with a robotic position matching task and reported many kinematic variables. The incidence of proprioception task failure where vision was occluded was 100% and 97% in UN+ participants in each study, compared to ∼60% and 58% of UN-participants. The presence of UN was also associated with increased performance variability and initial movement direction error.

### Functionally Defined UN

UN was assessed with a functional tool by Barra et al. [36] and Rousseaux et al. [42]. Both studies utilised the Catherine Bergego scale, and hence assessed all domains and modalities of UN. Similarly, both assessed proprioception with a representation task, and reported significant correlations between UN and longitudinal body axis deviation, and a significantly altered ability to judge body topography in those with UN compared to those without.

## Discussion

The primary objective of this review was to determine if the presence of UN is associated with more severe proprioceptive deficit in stroke affected populations. A total of 15 out of 19 study results supported this association. Three studies reported no association, two using a position matching task and one a body axis judgement to assess proprioception. Thus, the majority of studies in this review support the association between UN and impaired proprioception.

However, this conclusion is limited by the variety of assessments used and modest quality of many studies. Importantly, both proprioceptive impairment and UN have been assessed with multiple methods and assessment tools which precludes quantitative analysis and thus their collective interpretation. UN assessment often failed to account for the multimodal nature of the condition. The inconsistency in assessment found in the literature likely reflects the current state of clinical practice and forms the basis for the lack of evidence-based treatment options, which is of concern given the negative functional consequences associated with UN.

There were moderate issues with quality in most studies included in this review. Only one study justified their sample size, a useful indicator of pre-planning for a study [54]. In addition, non-consecutive recruitment methods were used in more than half of the studies. Importantly, none of the studies that used a convenience sample reported the characteristics of patients not eligible or refusing to be included in the study. Hence, it is difficult to generalise the results of this review to stroke populations given that the characteristics of participants that were *not* recruited into each study are unknown. These factors contribute to a considerable level of bias in the studies in this review. Taken together, the heterogeneity of both method and quality of the studies render it impossible to draw conclusions about the relationships between specific forms of UN, and different components of proprioception.

### Clinical Implications

#### Assessment of UN

The issue of inconsistent assessment of UN has previously been identified and is echoed in the results of this review [15, 22]. At present, there is only a single assessment tool that covers all three hemi-spaces, the CBS. The CBS is also the only published assessment tool that incorporates functional tasks of both the upper and lower limb, and thus can directly guide clinical treatment decisions and provide a measure of neglect in the context of patient activities of daily living [55]. Most studies in the present review assessed UN in a single hemi-space, most commonly via pen and paper cancellation tasks. These tasks have mixed reliability, their validity was only tested against other cancellation tasks, and have no reported responsiveness [15]. In addition, the attentional demands of these tasks may not be high enough to detect subtle signs of UN. They also may allow for compensation [56]. Importantly, only two studies [36, 42] used the CBS.

There are additional issues with the assessment methods in studies that evaluated more than one form of UN. Those that assessed representational UN, used a scene or figure copying task, neither of which are standardised testing protocols. Furthermore, those tests have shown to be insensitive, and have questionable validity [16]. The two studies [49, 50] that assessed UN using a behavioural task used the Behavioural Inattention Test. This test assesses exclusively the peri-personal space, and involves tasks restricted to the upper limb. Hence, it fails to provide a functional assessment of UN in all three hemi-spaces. It also excludes the assessment of the lower limb. Taken together, the BIT only partially assesses UN and thus requires clinicians to use other tools to capture the full spectrum of possible impairment, which is not ideal in common care contexts of high workload and reduced threshold of patient fatigue.

#### Assessment of Proprioception

There was considerable heterogeneity in the protocols used to assess proprioception in the included studies. Three different subtypes of higher-level tasks were assessed – body axis judgement, laterality, and body topography. Two different laterality judgement protocols were reported, one using pictures of hands and the other pictures of human stick figures. Three different measures of body topography were used, differing in orientation (first or third person) and nature (pictorial or actual size) of the presented body, and the motor task required (rearranging, pointing, or searching). In addition, three position matching protocols were used, two of which have not been validated. Finally, the assessment of force judgement was absent from the studies of this review. Because of the limited number of studies and the high variability in testing procedures, it was difficult to draw a strong conclusion regarding the impact of proprioceptive deficits in UN.

However, proprioceptive impairment, notably ‘high level’ impairment, is indeed implicated in UN. This is important given that typical clinical assessment of proprioception fails to capture multiple levels of the sense [for review see 57]. The standardised, clinically used tools to test proprioception include the Erasmus Modification of the Nottingham Sensory Assessment and the Rivermead Assessment of Somatosensory Perception. These tools are classified as ‘low-level’ proprioceptive assessments and use an ordinal grading system, defining the patient’s proprioception as either normal, impaired, or having “no proprioception at all” [58, 59]. Thus, it is impossible to make distinctions within the three grades and the grading system is not sensitive to small changes in proprioception. Furthermore, the correlation of ‘low-level’ proprioceptive tests to patient function and activity is low, or absent entirely [53]. Hence, clinical tools to measure proprioception fail to capture ‘high-level’ aspects of proprioception that are likely impaired in this population. There is a clear need for a standardised, clinically applicable test battery that encompasses multiple aspects of proprioception. This would allow clinicians to identify the full spectrum of proprioceptive impairment present in this, and other populations.

#### Study Limitations

The average size of the UN+ group in the studies included in this review was 13 (SD ±14, range 3-59) participants, and the maximum was 59. Thus, the present review is limited by the relatively small sample sizes of the studies, which compromises the generalization of our findings.

Another limitation in our study was the lack of success in obtaining some full text and additional data from authors. Seven abstracts screened were of unpublished studies. All authors were contacted for full text, however in all cases it was not forthcoming. Hence, all seven were excluded at full text review. A further eight studies were excluded at full text review due to insufficient reporting of data about UN+ and UN-groups. All authors were contacted to request data, but in all eight cases the data was either unavailable or no reply was received. Given the collective sample size of these studies (n=504), the inclusion of these data could change the strength of, or the findings themselves of the present review.

## Conclusions and Future Directions

This is the first systematic review that has summarised the evidence investigating proprioception in UN after stroke. There is moderate quality evidence that people with UN after stroke are more likely to have proprioceptive deficits than those without UN. These deficits occur across a variety of different subtypes of UN and levels of proprioception. A large limitation of the studies included in this review is that the assessment of both UN and proprioception is highly inconsistent, which likely reflects current clinical practice. Future investigations in this area should prioritise functional assessments of UN to provide evidence that can be easily translated to clinical practice. Investigation of proprioception via force judgement is absent from the literature in UN, and hence is an important area for future research. There is a clear need for a standardised, clinically applicable test battery that encompasses multiple aspects of proprioception.

## Supporting Information

**S1. Search strategy**

**S2. Reasons for full text exclusion.** 1. Abstract only, 2. Insufficient data, 3. No UN-group, 4. No UN+ group, 5. No proprioception measure, 6. Participants with previous stroke, 7. Ineligible study design

**S3. AXIS quality assessment study results**

**S4. PRISMA Checklist**

